# In-field metagenome and 16S rRNA gene amplicon nanopore sequencing robustly characterize glacier microbiota

**DOI:** 10.1101/073965

**Authors:** Arwyn Edwards, Aliyah R. Debbonaire, Samuel M. Nicholls, Sara M.E. Rassner, Birgit Sattler, Joseph M. Cook, Tom Davy, André Soares, Luis A.J. Mur, Andrew J. Hodson

## Abstract

In the field of observation, chance favours only the prepared mind (Pasteur). Impressive developments in genomics have led microbiology to its third “Golden Age”. However, conventional metagenomics strategies necessitate retrograde transfer of samples from extreme or remote environments for later analysis, rendering the powerful insights gained retrospective in nature, striking a contrast with Pasteur’s dictum. Here we implement highly portable USB-based nanopore DNA sequencing platforms coupled with field-adapted environmental DNA extraction, rapid sequence library generation and off-line analyses of shotgun metagenome and 16S ribosomal RNA gene amplicon profiles to characterize microbiota dwelling within cryoconite holes upon Svalbard glaciers, the Greenland Ice Sheet and the Austrian Alps. We show in-field nanopore sequencing of metagenomes captures taxonomic composition of supraglacial microbiota, while 16S rRNA Furthermore, comparison of nanopore data with prior 16S rRNA gene V1-V3 pyrosequencing from the same samples, demonstrates strong correlations between profiles obtained from nanopore sequencing and laboratory based sequencing approaches. gene amplicon sequencing resolves bacterial community responses to habitat changes. Finally, we demonstrate the fidelity and sensitivity of in-field sequencing by analysis of mock communities using field protocols. Ultimately, in-field sequencing potentiated by nanopore devices raises the prospect of enhanced agility in exploring Earth’s most remote microbiomes.

## INTRODUCTION

Microbes drive biogeochemical processes at all scales within the biosphere(Falkowski et al., 2008). The ubiquity of microbes within every manner of niche on Earth is underpinned by the expansive range of microbial genomic diversity, spanning all three domains(Hug et al., 2016). However, our view of microbial diversity on Earth is constantly changing as technological advances reveal novel groups and associations of microbes(Rinke et al., 2013; Hug et al., 2016)). Within recent decades, sequencing of DNA extracted from environmental matrices, either amplified(Schmidt et al., 1991) or sequenced directly via shotgun metagenomics(Tyson et al., 2004) makes the prospect of a predictive, multi-scaled understanding of microbial interactions with the Earth system increasingly tangible(Widder et al., 2016).

Typically, such investigations require the transfer of collected microbiota for nucleic acid extraction, sequencing and bioinformatic analysis at facilities with a high level of infrastructure. These strategies are well established. However, the necessity to transfer samples from remote locations to the laboratory incurs several disadvantages, including the potential loss or corruption of unique samples in transit as well as biases incurred by taphonomic degradation during storage and subsequent extraction(Klein, 2015). In particular, the delays incurred via this strategy means that the process of gaining genomic insights to microbial processes within the natural environment is divorced from the environment in which they occur, rendering the exploration of microbial diversity a reactive, rather than proactive activity.

Pocket sized, USB-driven, nanopore DNA sequencers (e.g. the Oxford Nanopore Technologies Ltd MinION) potentially offer a novel direction for the versatile characterization of microbiomes. Highly portable DNA sequencing strategies promise the generation and analysis of DNA sequences within field settings, thus removing the risk of sample degradation or loss in cold chain to a home laboratory and significantly accelerating the process of generating insights.

While MinION sequencing has been applied for in-field genomic epidemiology studies of the West Africa Ebola virus outbreak(Quick et al., 2016) and genome sequencing on the International Space Station(Castro-Wallace et al., 2017) its application to in-field microbial ecology applications has been limited. However, the potential for characterizing microbial communities via nanopore-based shotgun metagenomics or amplicon sequencing on the MinION platform has been established within laboratory-based sequencing experiments using contrived or well-known communities(Brown et al., 2017; Kerkhof et al., 2017). Astrobiologists have long used polar field sites as low-biomass environment analogues as a testbed for life detection strategies, and the incorporation of MinION in trials for life detection strategies offers some insight to the performance of MinION in remote and challenging environments. This includes a trial of the MinION platform’s technical performance and endurance in Antarctica (Johnson et al., 2017) and the incorporation of MinION-based metagenomics along with culturing as a proof-of-principle life detection system in the Canadian High Arctic(Goordial et al., 2017). This latter study demonstrated the potential utility of MinION sequencing in field settings but due to poor weather and limited internet connectivity was limited to only one successful experiment where data could be processed using a cloud-based bioinformatics platform.

Therefore, applications of nanopore sequencing in the field to date have typically required labour-intensive sequencing library preparation following protocols developed for laboratory use, and data analysis required base-calling and/or annotation using remote servers. Reliance on cold chain transfer of reagents and sequencing flow cells also presents an important limitation. However, the implementation of shotgun metagenomics and 16S rRNA gene amplicon analysis for in-field sequencing of complex microbiomes in conditions typical for microbial ecologists in remote environments requires these obstacles to be overcome.

In this study we show the applicability of nanopore sequencing for rapid taxonomic characterization and comparison of microbial communities in remote environments. Moreover, we benchmark the performance of in-field sequencing and bioinformatics approaches used by sequencing of mock communities analysed following in-field protocols and re-analysis of samples previously investigated by second-generation sequencing. For our analyses we selected cryoconite, a photo- and heterotrophic microbe-mineral aggregate darkening the ice surfaces of glaciers and ice sheets (Cook et al., 2015) as target microbiota. Cryoconite aggregates are home to a diverse range of microbial life, with estimated diversity and activity rates comparable to certain soils (Anesio et al., 2009; Cameron et al., 2012) in spite of its icy environs. We show shotgun metagenomic and 16S rRNA gene amplicon sequencing using rapid library preparation protocols and fast, laptop-based taxonomic classification tools are capable of robustly characterizing and comparing microbial communities while in the field. Our results raise the prospect of investigating Earth’s microbiomes at source.

## MATERIALS AND METHODS

### Sampling

Sampling and sequencing was performed in Svalbard and Greenland, while archived materials from three Svalbard glaciers and two alpine glaciers were used in the benchmarking experiments. In all cases, cryoconite samples were collected using a disinfected turkey baster which was used for the aspiration of debris to 15 mL sterile centrifuge tubes and transferred within hours on ice in ambient temperatures of +2-4°C to the field lab or camp where they were either extracted directly or stored in a −20°C freezer. While a common DNA extraction protocol is employed, over the course of this study, protocols and tools for nanopore sequencing have advanced rapidly and we describe the implementation of nanopore sequencing in remote locations with differing levels of infrastructure. Each experiment is therefore described in turn and summarized in Table 1.

**Table 1:** Summary of sequencing experiments.

| **Experiment** | **Analysis type** | **Study Site** | **DNA extraction** | **Library kit** |
| --- | --- | --- | --- | --- |
| Svalbard RAD001 | Shotgun | Foxfonna | Mod-Powersoil | RAD001 |
| Greenland RAD001 | Shotgun | Russell glacier | Mod-Powersoil | RAD001 |
| Austria RAD003 | Shotgun | Rotmoosferner | Mod-Powersoil | RAD003 |
| Austria LRK001 | Shotgun | Rotmoosferner | Mod-Powersoil | LRK001 |
| Mock LRK001 | Shotgun | Lab | N/A | LRK001 |
| Svalbard 16S | 16S rRNA gene | Vestre Brøggerbreen | Mod-Powersoil | RAB201 |
| Arctic-Alps 16S | 16S rRNA gene | Lab | Mod-Powersoil | RAB201 |
| Mock-Spike 16S | 16S rRNA gene | Lab | N/A | RAB201 |

### In-field DNA extraction

For cryoconite studies in the present study, community genomic DNA was extracted directly from cryoconite debris in field conditions. PowerSoil® DNA Isolation kits (MoBio, Inc.) are commonly used to extract high quality DNA from environmental matrices rich in nucleic acid processing enzyme inhibitors, including glacial samples (Edwards et al., 2011; Cameron et al., 2012; Musilova et al., 2015; Stibal et al., 2015; Lutz et al., 2016). We therefore opted to amend the PowerSoil® DNA isolation protocol for use within a minimalistic field laboratory to generate DNA extracts compatible with nanopore sequencing protocols.

Generally, the protocol was implemented as per the manufacturer’s instructions with the following variations: The starting material was increased from 0.25 g to 0.27-0.29 g wet weight to account for moisture content prior to bead-beating within PowerBead tubes. Bead-beating was conducted using a Vortex Genie 2 (Scientific Industries, Inc) fitted with a MoBio (MoBio, Inc.) tube adapter or a IKA Works, Inc MS2 S8 Minishaker. Since the PowerSoil® protocol typically uses benchtop centrifuges capable of 15-17,000 ×g, the use of a Gilson, Ltd GmCLab Microcentrifuge capable of generating <2,900 ×g required elongated spin times of four minutes for protein precipitation following the addition of solution C2 and inhibitor removal following the addition of solution C3. For all other centrifugation steps, spin times of one minute were sufficient. To increase DNA concentration, DNA was eluted in 70 µL solution C6, after a five minute incubation at room temperature. Two microlitre aliquots of each extract were than immediately quantified using Qubit 2.0 Fluorimeter dsDNA High Sensitivity assays (Invitrogen, Ltd) prior to sequencing. In the case of Greenland metagenomics, fluorimetry was performed with the Qubit device powered using a portable power-pack (PowerAdd, Inc.)

### Shotgun metagenomics

#### Svalbard: shotgun metagenomics experiment, Foxfonna ice cap

On 18 August 2016, cryoconite debris from four cryoconite holes was collected as described above on the un-named outlet glacier of Foxfonna ice cap (78° 08’N, 16° 07’ E; (Rutter et al., 2011; Gokul et al., 2016).) Cryoconite was thawed the following day and DNA extracted as described. The analyses were performed in a room without prior laboratory infrastructure. Equimolar pools of cryoconite DNA provided 185 ng DNA used for a transposase-based 1D rapid sequencing library preparation exactly as specified by Oxford Nanopore Technologies, Ltd (Rapid Sequencing of genomic DNA for the minION device using SQK-RAD001) using a PCR cycler for the 30°C and 75°C incubation steps in a Hybaid Omni-E PCR cycler.

1D sequencing was directly performed using MinION Mk 1 device containing a R9 flow cell (Oxford Nanopore Technologies, Ltd) loaded conventionally in 150 µL sequencing mix. The MinION Mk 1 device was controlled using MinKNOW 1.0.5 (Oxford Nanopore Technologies, Ltd). Sequencing and read processing were perfomed on a Lenovo® ThinkPad® X1 Carbon running Windows 10 with Intel® Core™ i7-5600U Processor, 8GB RAM and a 512GB solid state drive. 1D basecalling was performed remotely via the Oxford Nanopore Technologies, Ltd. Metrichor platform. Template strand nanopore reads in .fast5 format were converted to fasta using the pore_parallel GUI of poRe 0.17 (Watson et al., 2015) running in R 3.3.1.

#### Greenland Ice Sheet margin: shotgun metagenomics experiment, Russell glacier

On 16 June 2017, cryoconite debris from four cryoconite holes was collected as described from four cryoconite holes on Russell glacier at the south western margin of the Greenland Ice Sheet (N67° 8’08.871’ 05 ° 08’05.016′ W(Edwards et al., 2014b)) DNA extraction and sequencing was performed in a field camp tent with AC generator power for DNA extraction only. Equimolar pools of cryoconite DNA provided 185 ng DNA used for transposase based 1D rapid sequencing library preparation as specified by Oxford Nanopore Technologies, Ltd (Rapid Sequencing of genomic DNA for the minION device using SQK-RAD002) but holding the library tube in gloved hands for the 30°C incubation step and immersion in hot water stored in an insulated mug for 75°C inactivation.

The library (75 µL) was loaded in the SpotON port of a R9.4 flow cell (Oxford Nanopore Technologies, Ltd) run using a MinION MK1 device. Due to the cold (<0°C) ambient temperatures, to maintain optimal sequencing temperature, the MinION device was enclosed in an insulated cool bag during sequencing (Tesco, Ltd). Sequencing and read processing were perfomed on a Dell XPS15 laptop running Windows 10 Intel® Core™ i7-6700HQ Processor, 32GB RAM and a 1TB solid state drive, running on internal battery. Sequencing was controlled using MinKNOW 1.6.11 (modified for offline use, courtesy Oxford Nanopore Technologies, Ltd) and 1D basecalling performed locally with Albacore v1.1.0 (Oxford Nanopore Technologies, Ltd).

#### Benchmarking: shotgun metagenomics experiment, Rotmoosferner, Austrian Alps

We sought to compare the performance of in-field nanopore metagenome sequencing with established strategies for metagenome sequencing and compare the performance of rapid library kits based upon aqueous reagents used in the field study with the current state of the art, namely rapid library kits based upon freeze-dried reagents.

Therefore, we conducted nanopore sequencing of samples with corresponding, publicly available metagenome data. Samples RF1 and RF6, from alpine cryoconite were collected from the surface of Rotmoosferner glacier in the Austrian Alps in September 2010. A shotgun metagenomic sequencing library of the same two samples in an equimolar pool with 12 other cryoconite community genomic DNA samples from Rotmoosferner was analysed by Illumina sequencing previously (Edwards et al., 2013b) and is available at MG-RAST (4 491 734.3).

Community genomic DNA was extracted in 2017 from 250 mg cryoconite sediment from samples RF1 and RF6 stored at −80°C using the field-modified PowerSoil method in our home laboratory. To generate rapid 1D libraries, 400 ng DNA from the samples were used in transposase-based protocol as specified by Oxford Nanopore Technologies, Ltd (Rapid Sequencing of genomic DNA for the minION device using SQK-RAD003) with incubation steps in a PCR Cycler (G-Storm Direct, Ltd). Library was loaded and sequenced on a R9.4 flow cell as described above for the *Greenland* experiment, using MinKNOW 1.7 and Albacore v2.02 for sequencing and basecalling respectively.

Storage and transportation of molecular biology reagents and sequencing flow cells at cold temperatures represents a critical limitation to the utility of nanopore sequencing for characterizing microbial communities in remote locations. We therefore tested lyophilized versions of the rapid library kit (SQK-LRK001) and ambient-shipped R9.4 flow cells provided by Oxford Nanopore Technologies Ltd. in a repetition of the above experiment. The experiment was performed without laboratory resources. Briefly, 400 ng of community DNA in a 10 µL volume was used to suspend lyophilized fragmentation mix, with incubation at room temperature with inactivation and immersion in hot water stored in an insulated mug for 80°C inactivation for one minute each. The resulting solution was incubated with rapid adapter for five minutes before gentle mixing with 65 µL priming buffer and flow cell loading before sequencing as described above.

#### Benchmarking: sequencing of mock community DNA using lyophilized rapid kit

Finally, to establish the fidelity with which in-field DNA sequencing can capture microbial diversity, we performed DNA sequencing on the ZymoBIOMICS^™^ Microbial Community standard (provided as DNA, D3606, Zymo Research, Inc.) using the lyophilised rapid kit and field protocols, albeit in a laboratory setting.

The ZymoBIOMICS^™^ Microbial Community standard comprises a mixture of eight bacterial species (*Pseudomonas aeruginosa, Escherichia coli, Salmonella enterica, Lactobacillus fermentum, Enterococcus faecalis, Staphylococcus aureus, Listeria monocytogenes, Bacillus subtilis*) each comprising 12% of the DNA by mass, and two yeast species (*Saccharomyces cerevisiae* and *Crypotoccus neoformans*) each comprising 2% of the DNA by mass. The manufacturers state that the mixture contains <0.01% foreign microbial DNA and the composition is stable to within a relative abundance deviation of 15% on average. Further details on the ZymoBIOMICS^™^ Microbial Community standard are available at: https://www.zymoresearch.eu/media/amasty/amfile/attach/_D6305_D6306_ZymoBIOMICS_Microbial_Community_DNA_Standard_v1.1.3.pdf

Briefly, 400 ng of the ZymoBIOMICS^™^ Microbial Community standard in a total volume of 10 µL was used, exactly as described above for lyophilised rapid kit sequencing.

#### Bioinformatics for shotgun metagenomics

For all experiments, sequencing reads were analysed using Kaiju(Menzel et al., 2016), a taxonomic classifier which is offered as a webserver (http://kaiju.binf.ku.dk.) or as a standalone version. Kaiju seeks protein-level matches in all possible reading frames using the Burrows-Wheeler transform and has been demonstrated to be rapid yet sensitive (Menzel et al., 2016). We report Kaiju analyses conducted using the NCBI *nr*+*euk* database (version: 05 May 2017) containing 103 million protein sequences from microbes, viruses and selected microbial eukaryotes with low complexity filtering and in *greedy* mode (Minimum match: 11; minimum match score: 75; allowed mismatches: 5). Nanopore sequencing data from in-field metagenomes are available at the European Nucleotide Archive (ENA) accessions ERR2264275 - ERR2264278 while mock community metagenomes are available at ENA PRJEB30868.

### 16S rRNA gene sequencing

#### Svalbard: 16S ribosomal RNA gene sequencing, Vestre Brøggerbreen

On 10 July 2017, cryoconite debris was collected from six cryoconite holes on Vestre Brøggerbreen (78° 54’7 N, 011°43’8 E) on Svalbard. Three cryoconite holes (VB1-3) were covered by snow and superimposed ice while three cryoconite holes (VB4-6) were exposed. To test the hypothesis that the presence or absence of snow/ice cover incurs changes in the bacterial community structure, 16S ribosomal RNA gene sequencing was conducted. Samples were transferred to the NERC Arctic Research Station Ny Ålesund within three hours and DNA extracted and quantified as described above within the field lab.

Bacterial 16S rRNA genes were amplified from 50 ng DNA per sample diluted to 10 µL nuclease free water in 50 µL with 1 × LongAmp Taq master mix (New England Biolabs, Inc), 1 µL 16S rRNA gene barcoded primer (ONT-SQK-RAB-201, Oxford Nanopore Technologies, Ltd) and 14 µL nuclease free water. Each sample was allocated to a barcoded primer. A no template control (nuclease free water) and extract control (blank extract) were amplified and sequenced in parallel. PCR was conducted as specified in the Oxford Nanopore Technologies protocol for rapid, barcoded sequencing of 16S rRNA genes (ONT-SQK-RAB-201, Oxford Nanopore Technologies, Ltd) with the modification that 30 cycles of PCR were performed. PCR was performed using a 8 well MiniPCR device (Amplyus, Inc.) controlled by a Windows laptop. The resulting PCR products were purified using magnetic beads (Ampure XP, Beckman Coulter, Ltd) and eluted in 10 µL buffer (10 mM Tris, 50 mM NaCl, pH 8.0) prior to Qubit quantification. Equimolar quantities of each barcode were pooled and sequenced using a R9.4 flow cell as described above for the *Greenland* experiment prior to 1D basecalling and barcode demultiplexing in Albacore v1.10.

#### Benchmarking: nanopore 16S rRNA gene discrimination of Arctic and alpine cryoconite

To compare the performance of in-field nanopore 16S rRNA gene sequencing with established strategies for 16S rRNA gene profiling, we conducted nanopore sequencing of cryoconite samples with corresponding, publicly available 16S rRNA gene data spanning Svalbard and Austrian glaciers. V1-V3 16S rRNA amplicon pyrosequencing data for these samples are available at available at EBI-SRA (PRJEB5067-ERP004426). Details of sample collection, pyrosequencing and data processing are described elsewhere (Edwards et al., 2014b).

Ten samples (Svalbard: three pairs of samples from Austre Brøggerbreen [AB], Midtre Lovénbreen [ML] and Vestre Brøggerbreen [VB]. Austria; two pairs of samples from Gaisbergferner[GB] and Rotmoosferner[RM]) archived at −80°C since collection were used to generate fresh DNA extracts as described above. Bacterial 16S rRNA genes were amplified and sequenced exactly as described for the Svalbard 16S ribosomal RNA gene sequencing experiment, with the exception that a 96 well PCR cycler (G-Storm Direct, Ltd) was used for amplification and Albacore v2.02 used for basecalling and demultiplexing by barcode. Samples were assigned an individual barcode primer, with co-amplification and sequencing of no-template and extract controls.

#### Benchmarking: 16S rRNA gene nanopore sequencing of mock communities

To evaluate the accuracy and sensitivity of 16S rRNA gene nanopore sequencing we analysed mock communities. The ZymoBIOMICS^™^ Microbial Community Standard was used to represent a baseline community of eight members with 16S rRNA genes. To establish the accuracy of taxonomic assignment using uncorrected 16S rRNA gene reads and the sensitivity of our protocol to be able to detect taxa present above and below 1% theoretical relative abundance, we spiked the ZymoBIOMICS^™^ Microbial Community Standard with genomic DNA of *Micrococcus luteus* NCTC2665, an actinobacterial strain with a 2.65Mbp genome and only one functional 16S rRNA gene (Young et al. 2010). *M*. *luteus* DNA was extracted and quantified as above, from an overnight nutrient broth culture stored on ice for one hour prior to extraction. Genomic DNA from *M*. *luteus* was spiked in to create mixtures where the theoretical relative abundance of the *M*. *luteus* 16S rRNA gene was 0%, 0.4%, 0.8% and 1.2% of the total bacterial 16S rRNA genes. Otherwise, amplification and sequencing of 16S rRNA genes were conducted exactly as described above for the comparison of Arctic and alpine cryoconite with the exception that negative and extraction controls were not sequenced. It should be noted that two members of the community, *Escherichia coli* and *Salmonella enterica* share highly conserved (typically well above 97% identity, e.g. Fukushima et al. 2002) 16S rRNA genes therefore these taxa were treated as a single entity in data processing.

#### 16S rRNA gene data analysis

Reads were base-called and de-multiplexed using Albacore v1.10 and v2.02 for implementation and benchmarking experiments, which were converted to fasta for taxonomy assignment. In developing a strategy for nanopore 16S rRNA amplicon analysis, our primary considerations were to leverage the short turnaround time and in-field flexibility of the MinION platform while mitigating the impacts of relatively high error rate per base. We therefore opted to directly assign higher-level taxonomy to Albacore demultiplexed reads via the SINTAX taxonomic classifier in usearch v10.0.240(Edgar, 2016) against taxa in a species-level identity curated database of Ribosomal Database Project version 16 taxa. A confidence level of 0.75 was chosen for each taxonomic assignment. The taxonomy assignment is available as supplementary data.

The numbers of reads assigned per taxon were counted in MS Excel and the relative abundance of reads per taxon used for separate downstream analysis. Multivariate analyses of bacterial community structure using Bray-Curtis distances of fourth root transformed taxon relative abundance data were performed in Primer-6.1.12 & Permanova+1.0.2 (Primer-E, Ltd, Ivybridge, UK). Principal Coordinates Analysis (PCoA) and group-average Hierarchical Cluster Analysis (HCA) were used as unsupervised data exploration, while hypotheses were tested using PERMANOVA with unrestricted permutation of raw data and 9,999 ordinations. Reads from the implementation, comparison and mock community experiments are available from ENA at ERR2264279 - ERR2264283, ERR2264284 - ERR2264293 and PRJEB30868.

## RESULTS AND DISCUSSION

We report the successful implementation of nanopore sequencing for generating taxonomic profiles of microbiota in field environments, both from shallow-depth shotgun metagenome and targeted, 16S rRNA gene amplicon sequencing approaches. Our analyses reveal nanopore sequencing generates taxonomic profiles coherent with conventional workflows in molecular microbial ecology. These results raise the prospect that highly portable DNA sequencing can be applied to characterise microbiomes rapidly while researchers are deployed in remote field sites and thus help inform experimental or survey planning and analysis within the field.

### In field metagenomics of Arctic cryoconite

Here, we report the generation of two metagenomes from Svalbard and Greenland cryoconite by in-field nanopore sequencing, and the taxonomic classification of microbial diversity (FIG 1).

**Figure 1:**
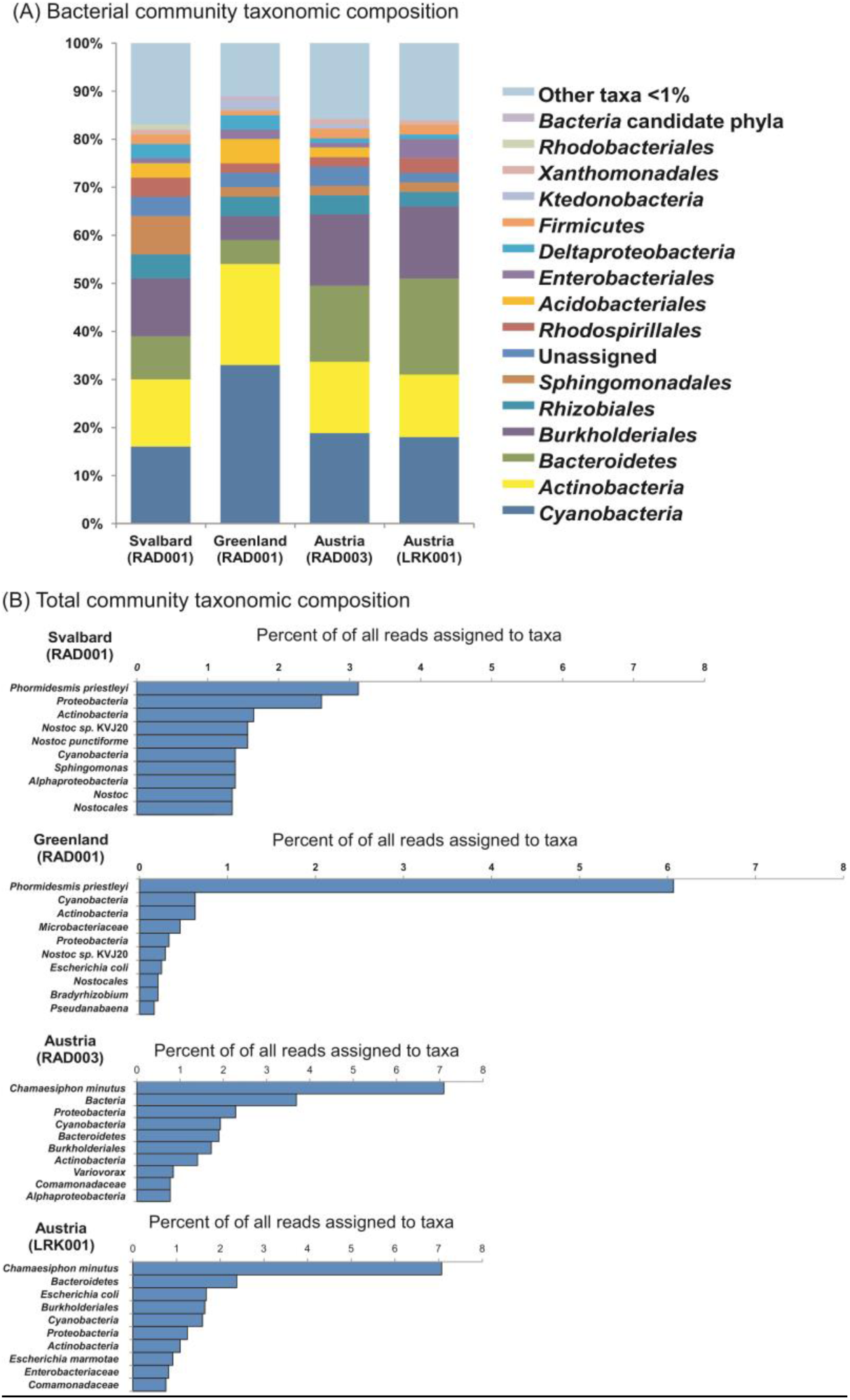
In field metagenome sequencing reveals taxonomic distribution of cryoconite microbiota. (A): shows bacterial taxa present at abundance greater than 1% of the bacterial community while (B) shows the top ten taxa as a percentage of reads assigned to any taxon. Svalbard and Greenland datasets were generated in the field, while Austria datasets were generated using field protocols to compare aqueous (RAD003) and lyophilised library preparation (LRK001) approaches.

For the *Svalbard* metagenomics experiment, community DNA was extracted from Foxfonna cryoconite and sequenced using rapid library preparation, with 3,514 reads successfully basecalled. Of these, through analysis using the Kaiju classifier, 2,305 reads could be assigned to named taxa, of which 2,265 reads were assigned as bacterial in origin. Consistent with earlier V1-V3 region 16S rRNA gene amplicon sequencing of cryoconite from Foxfonna ice cap(Gokul et al., 2016) the community was dominated by *Proteobacteria* (*Alphaproteobacteria* and *Betaproteobacteria*) followed by *Actinobacteria, Cyanobacteria* and *Bacteroidetes* (FIG 1). Importantly, the most abundant species-level match was to *Phormidesmis priestleyi*, representing 3% of the reads assigned to any taxon. The cyanobacterium *Phormidesmis priestleyi* is frequently detected in Arctic cryoconite and is thought to act as an ecosystem engineer of cryoconite granules, with its cyanobacterial filaments entangling inorganic debris to form darkened microbe-mineral aggregates on the ice surface (Langford et al., 2010; Cook et al., 2016; Gokul et al., 2016; Uetake et al., 2016).

The *Svalbard* metagenomics experiment represents an initial use of rapid library preparations to generate taxonomic profiles in field environments, with base-calling and subsequent analysis performed using online resources. Additionally, this experiment represents sample-to-preprint communication of its initial analyses within 23 days of sample collection (Edwards et al., 2016). Considering the doubling times of Svalbard cryoconite sedimentary bacterial communities are in the same order (Anesio et al., 2010) this represents the metagenomic characterization of a microbial community within its generation time.

The reliance of our *Svalbard* metagenomics experiment described above on internet access and mains electricity represents an important limitation. We therefore sought to conduct a second experiment based in a typical field camp setting. Our *Greenland* metagenome sequencing experiment entailed collection of cryoconite from the margin of the Greenland Ice Sheet and its sequencing. DNA extraction was performed with access to a portable generator, but generator failure required all subsequent steps to be performed on battery power. Quantification powered by a portable power-pack while library preparation was performed without power, and DNA sequencing was performed on laptop batteries. Freezing air temperatures presented challenges for maintaining liquid reagents and optimal MinION temperatures, resolved by storing reagents warmed by body warmth in a down jacket, and operating the MinION in an insulated shield. These limitations constrained the endurance of the sequencing experiment, but of 2,372 reads sequenced, 796 reads were assigned to taxonomy and 692 reads matched bacterial taxa (FIG 1). The community revealed was dominated by *Cyanobacteria, Proteobacteria* (*Alphaproteobacteria, Betaproteobacteria, Gammaproteobacteria*) and *Actinobacteria*, again consistent with sequence data from Greenlandic cryoconite(Edwards et al., 2014b; Musilova et al., 2015; Cook et al., 2016). The most abundant species within the profile, *Phormidesmis priestleyi*, represented 6% of the reads assigned to any taxon. Our *Greenland* metagenome experiment therefore represents the in-field metagenomic sequencing of a microbial community in the resource-limited settings typical of Arctic field camps.

### Benchmarking in-field metagenomics: the utility of freeze-dried reagents for sequencing

An important limitation to all above experiments is the necessity of maintaining a cold-chain for sequencing reagents and nanopore flow cells. This presents logistical challenges for characterizing microbial communities in highly remote locations. We therefore tested an early-release lyophilized field kit provided by Oxford Nanopore Technologies, Ltd, now available as SQK-LRK001. We compared its performance to cold-chain dependent aqueous-based reagents (SQK-RAD003) for rapid generation of sequencing libraries. To do so, we re-sequenced cryoconite DNA samples from Rotmoosferner, an alpine glacier, which had previously been characterized using Illumina sequencing(Edwards et al., 2013a). Two experiments were performed. The first experiment was performed using field protocols implemented in a laboratory setting while the second experiment used reagents and R9.4 flow cells adapted for ambient shipping and storage and stored at room temperature for at least five days. Kaiju classified 18,694 out of 31,538 reads generated using the lyophilised reagent kit, of which 18008 were assigned to the bacterial domain (FIG 1) while 2847 out of 6844 reads generated by the aqueous kit were classified. Comparison of the aqueous (Austria-RAD003) and lyophilised (LRK001) libraries (FIG 1) from the same samples showed highly coherent taxonomic profiles at higher grade taxon (Pearson *r=*0.97, p<0.001) and species levels (FIG1A, FIG 1B). The Kaiju-profiled communities were dominated by *Proteobacteria* (*Betaproteobacteria, Alphaproteobacteria, Gammaproteobacteria*) with *Burkholderiales* prominent within the community, consistent with previously reported Illumina-based shotgun metagenomics analyses of the same samples (Edwards et al., 2013a). *Cyanobacteria* were dominated by *Chamaesiphon minutus* matching reads at 7% of all reads sequenced in both samples. *Chamaesiphon* sp. are a prevalent cyanobacterial species within mountain glacier cryoconite (Segawa et al. 2017) therefore the detection of *C*. *minutus* as the dominant taxon within the alpine metagenomic dataset is likely valid.

### Benchmarking in-field metagenomics: sequencing of mock communities

To provide an estimation of the fidelity and reproducibility of in-field shotgun metagneomics we sequenced the ZymoBIOMICS^™^ Microbial Community Standard using the lyophilised field kit (LRK001) in three sequencing experiments (FIG 2). For the three experiments 94% of 29,399 reads, 95% of 52,000 reads and 94% of 29,084 reads were respectively classified by kaiju. The notably higher percentage of reads classified compared to experiments with environmental communities likely represents the stronger representation of the well-known taxa in the mock community within sequence databases compared to taxa from poorly described environments. All the expected species from the mock community were detected in the sequencing runs at family level. Within the *Enterobacteriaceae* reads were assigned to the genera *Escherichia* and *Salmonella* which could be expected within the community, but also the closely-related genus *Shigella* at a relative abundance of 1%. The taxonomic composition of the communities was highly consistent between sequencing experiments (FIG 2A), and correlated well with the expected relative abundances (FIG 2B, Pearson *r=*0.67, *p=*0.02).

**Figure 2:**
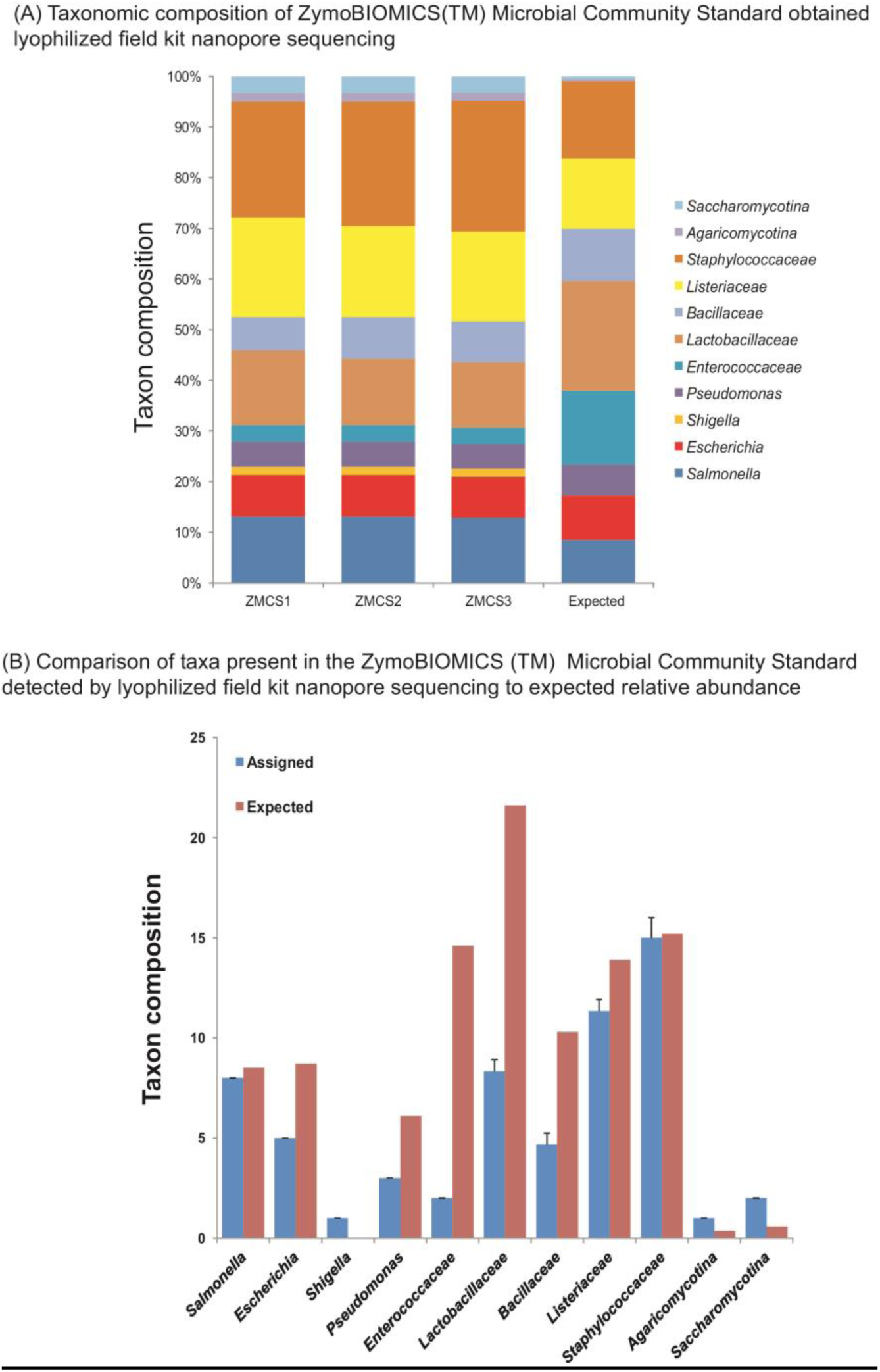
Benchmarking of in-field nanopore sequencing using lyophilized reagents. The ZymoBIOMICS Microbial Community Standard was sequenced in triplicate (ZMCS1-3) to compare against its expected composition and relative abundance profile. (A) shows the overall profile of all taxa detected at >1% relative abundance while (B) compares the relative abundance of assigned taxa against the expected composition.

The performance of Kaiju with nanopore data in our experiments matches the expectations of performance for Illumina data as described by its authors(Menzel et al., 2016). Since Kaiju uses protein-level matches to all possible reading frames it likely represents an effective tool for the rapid characterization of the taxonomic composition and implicit functional potential of microbial communities analysed by rapid library in-field nanopore sequencing of metagenomic DNA.

The performance of lyophilized field kit and ambient flow cells used for sequencing natural and mock communities using field protocols described above raises the prospect of metagenomic characterization of microbial communities in highly resource limited, remote environments.

### In field 16S rRNA gene sequencing

Amplicon sequencing of 16S rRNA genes represents a backbone technique in microbial ecology for the culture-independent investigation of microbial diversity (eg(Thompson et al., 2017)). In contrast with short read second-generation high throughput sequencing techniques, full-length 16S rRNA gene sequencing is possible with using nanopore(Li et al., 2016; Kerkhof et al., 2017) in laboratory settings. Therefore, we anticipated MinION based in-field sequencing of barcoded 16S rRNA genes could permit the semi-quantitative comparison of bacterial communities while in the field.

To test the feasibility of using barcoded 16S rRNA gene MinION sequencing to compare bacterial communities we analysed cryoconite holes which were either open to the atmosphere (VB1-3; n=3 FIG3 B) or covered by a layer of snow and superimposed ice (VB4-6; n=3 FIG3 B). On Arctic glaciers, cryoconite holes are seasonally open to the atmosphere in summer, permitting photosynthesis as a dominant carbon fixation route and supporting a diverse microbial community (Cook et al. 2015) Little is known of the dynamics of cryoconite communities before the seasonal recession of snow cover exposes the cryoconite hole, but it is likely the snow cover attenuates the flux of photosynthetically available radiation. We sought to test the hypothesis that the presence or absence of snow/ice cover incurs changes in the bacterial community structure.

**Figure 3:**
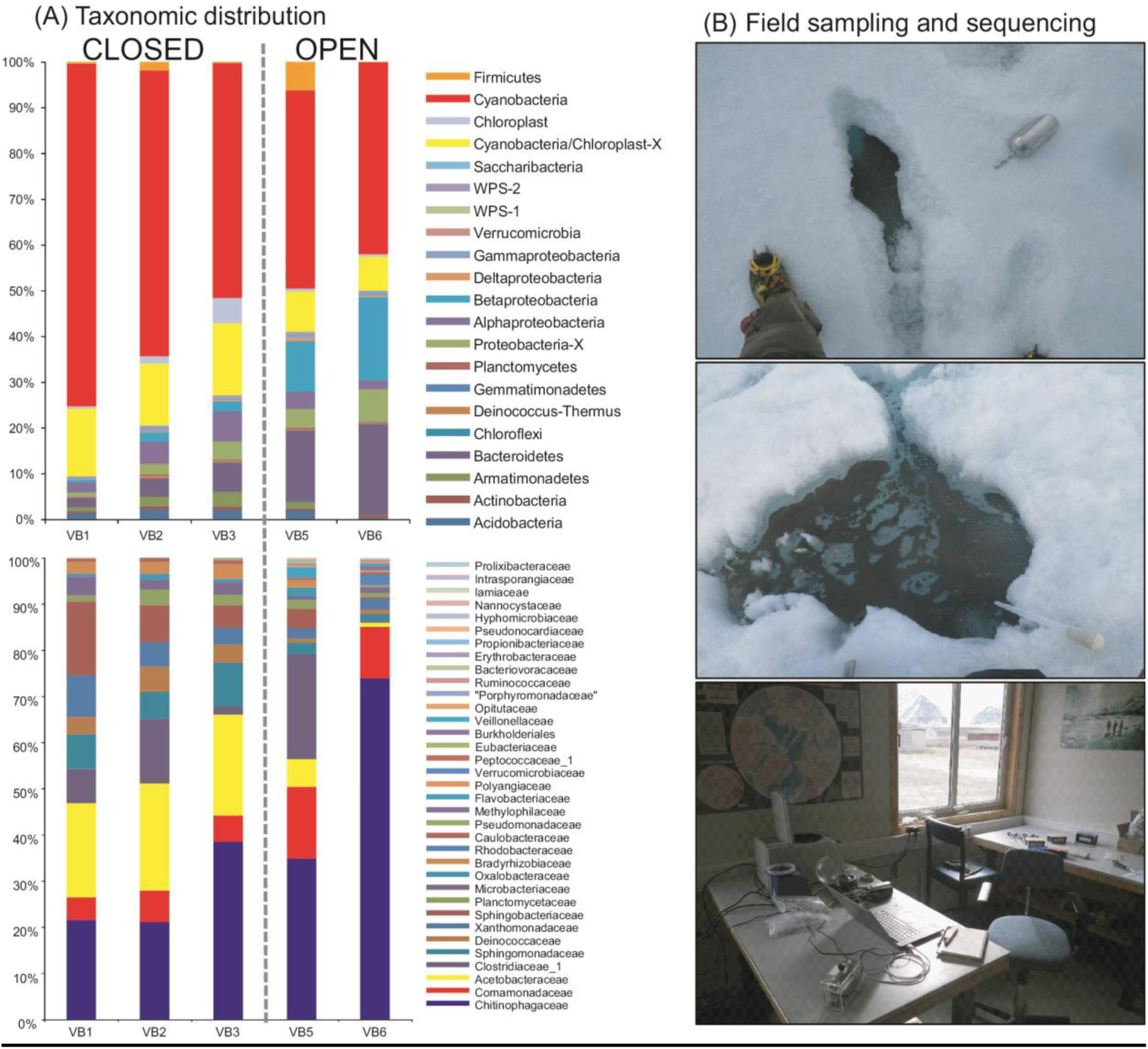
In-field 16S rRNA gene amplicon sequencing of cryoconite bacterial communities showing (A) phylum and family level taxonomic distribution of (B) open and closed cryoconite holes sampled on Vestre Brøggerbreen. The top image shows a “closed” cryoconite hole where the snow and superimposed ice cover has been displaced to reveal the hole, while the middle image shows a seasonally-open cryoconite hole. The bottom image shows the arrangement of equipment for DNA extraction, 16S rRNA gene PCR and nanopore sequencing in a field lab.

We amplified and co-sequenced a blank extraction control and a blank PCR control to detect the potential impacts of contamination arising from conducting 16S rRNA gene PCR (Salter et al., 2014), as preparing amplifications in a field laboratory setting may pose additional risks of contamination. A limitation of our experiment was the number of samples which could be processed in a single run, given the capacity of the eight-well PCR portable PCR cycler used. DNA was extracted and amplified from six samples, but PCR product from one sample (VB4, open) was lost during bead clean-up, resulting in sequencing failure for that sample.

Following basecalling and demultiplexing, 20,000 reads per sample were subjected to taxonomic analysis. Only seven sequences were returned for each of the negative controls, indicating the likely minimal impact of contamination in the field lab setting.

Although error-correcting approaches are in development for 16S rRNA gene nanopore sequencing(Li et al., 2016; Calus et al., 2018), considering the goal of preliminary characterization based upon rapid protocols compatible with field use, we opted to proceed with uncorrected reads, directly assigning taxonomy to each read and treating the cumulative abundance of reads matching discrete higher-level taxonomic affiliations as phylotypes.

Using the SINTAX classifier algorithm (Edgar, 2016) trained on a highly curated version of the RDP database, reads were classified within ca. 10 minutes on a laptop computer without the need for internet access. At a confidence level of 0.75, 15,643 reads per barcode were assigned to bacterial taxa on average (range: 13,017-17,593, 1 SD = 1725 reads). All community profiles were strongly dominated by *Cyanobacteria* at the phylum level with prominent contributions from *Proteobacteria* (*Betaproteobacteria* and *Alphaproteobacteria*) and *Bacteroidetes* broadly consistent with prior amplicon studies of Svalbard cryoconite (FIG 3). At both the phylum and family level, cryoconite communities ordinated and clustered clearly according to the open or closed status of the cryoconite hole (principal coordinates and hierarchical cluster analysis of fourth root transformed Bray-Curtis distances, FIG 4 A-D). At the family level, significant differences were apparent in the fourth root transformed Bray Curtis distances of taxon relative abundance (PERMANOVA: *F*=6.5, *p*(MC)=0.02). Our analyses are consistent with an influence of snow/ice cover on total bacterial community structure within cryoconite aggregates, however the increased abundance of cyanobacteria in the closed cryoconite holes is intriguing; it may be that the lower light levels in holes with thin snow/ice cover compared to open holes mitigate photoinhibitory effects (Cook et al. 2015) presenting a hypothesis which could be investigated further as a result of in-field sequencing. Our results underscore the potential for in-field 16S rRNA gene sequencing to inform investigations while deployed in remote environments.

**Figure 4:**
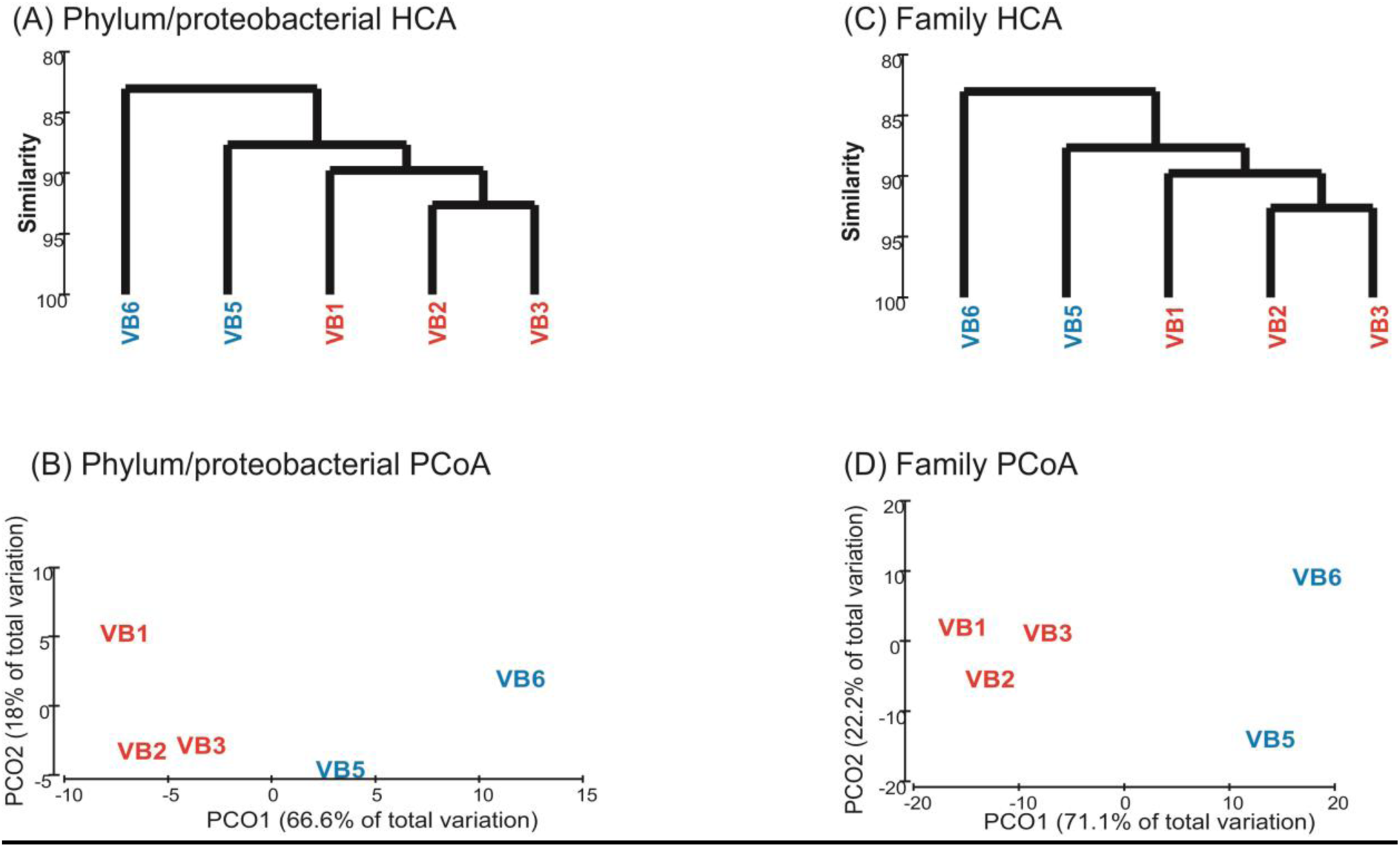
multivariate discrimination of cryoconite bacterial communities revealed by in-field 16S rRNA gene amplicon sequencing. Analyses are performed with data aggregated to phylum/proteobacterial class (A-B) or family-level taxa (C-D) with Hierarchical Cluster Analysis (HCA, subpanels A, C,) or Principal Cooordinates Analysis (PCoA, B,D,). Closed holes (VB1-3) and open holes (VB5-6) are ordinated by multivariate analysis of of fourth-root transformed Bray-Curtis distances of phylotype relative abundances.

### Benchmarking in-field 16S rRNA gene sequencing

To establish whether in-field 16S rRNA gene sequencing implemented as above offers results coherent with established, second-generation sequencing approaches we implemented our nanopore protocol on DNA samples which had previously been characterized by V1-V3 16S rRNA gene pyrosequencing(Edwards et al., 2014b). Our sample set comprised ten cryoconite samples from three Svalbard glaciers (n=2 each) and two alpine glaciers (n=2 each). Negative extraction and PCR controls as above produced one and four reads respectively. From 24,000 demultiplexed reads per sample, on average 20,882 reads were assigned to bacterial taxonomy (range: 9,021-24,000, 1 SD =6,518, FIG 5). *Cyanobacteria* with prominent contributions from *Proteobacteria* (*Betaproteobacteria* and *Alphaproteobacteria*) and *Bacteroidetes* dominated the phylum level taxonomic distributions of the community. At the phylum level, significant differences were observed between Arctic and alpine cryoconite communities (PERMANOVA; *F*=2.8866, *p*=0.02) with discrete ordination apparent (principal coordinates analysis of Bray-Curtis distances, FIG 5). At the improved taxonomic resolution of family level assignments, the effect of parent glacier is highly significant (PERMANOVA; *F*=5.032, *p*=0.001). In this regard, the outcomes of nanopore-based 16S rRNA gene sequencing are highly coherent with prior pyrosequencing analyses(Edwards et al., 2014b).

**Figure 5:**
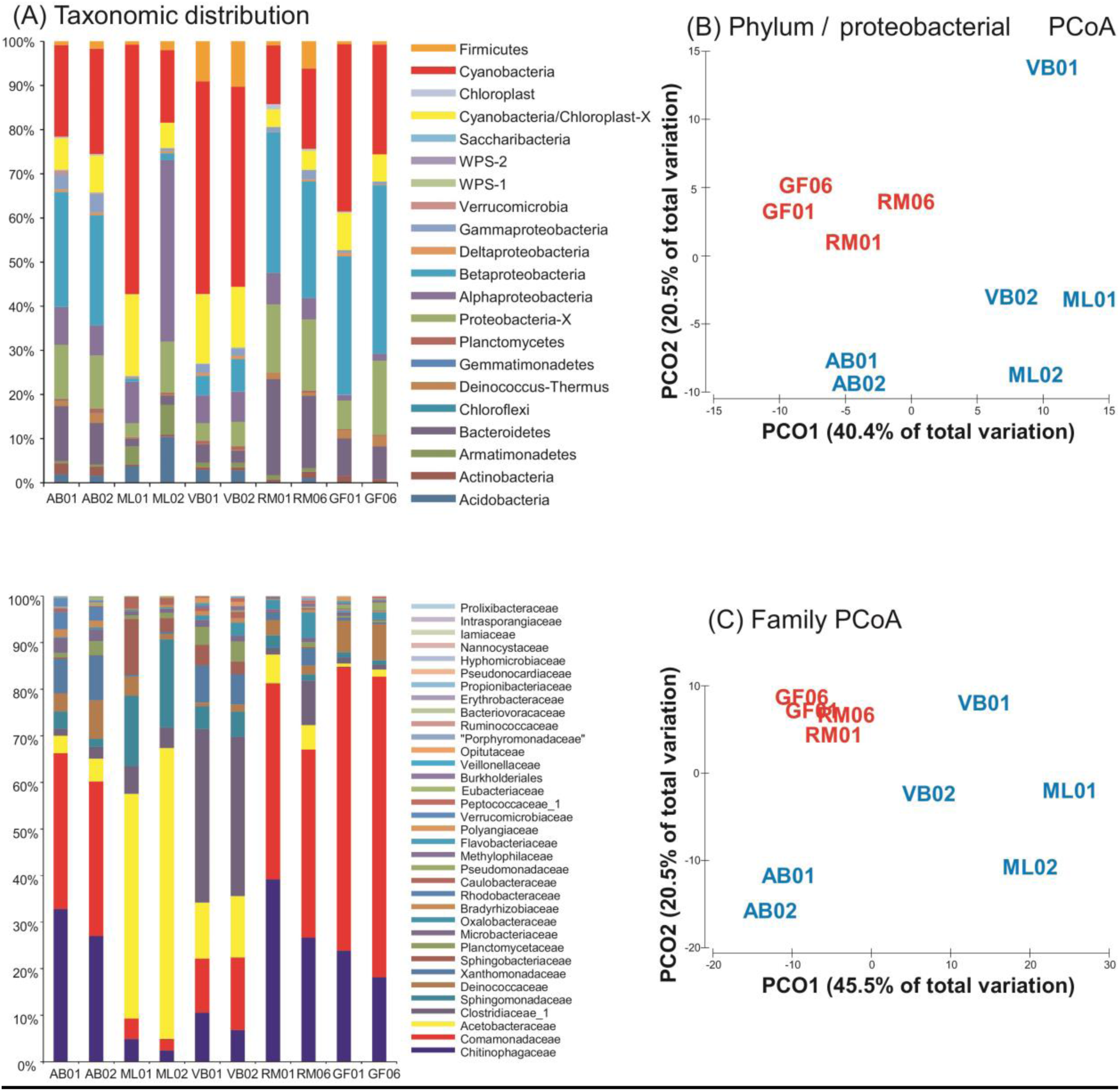
Benchmarking nanopore 16S rRNA gene amplicon sequencing of cryoconite bacterial communities by comparison with laboratory-generated data. (A) Phylum and family level taxonomic distribution of cryoconite bacterial communities and (B) phylum and (C) family level taxon distributions used for principal coordinates analysis of fourth-root transformed Bray-Curtis distances of phylotype relative abundances discriminate between Arctic and alpine cryoconite communities. Arctic glaciers (blue, AB, ML,VB) and Alpine glaciers (red, GF, RM) are clearly ordinated.

To further compare the assignment of taxa following nanopore-based 16S rRNA gene sequencing with the pyrosequencing dataset of the same samples, we correlated the log relative abundance of dominant taxonomic groups (FIG 6). Significant, strongly positive Pearson correlations between the log relative abundances of key taxonomic groups were apparent between nanopore and pyrosequencing data: *Acidobacteria* (*r*=0.82, *p*=0.004), *Alphaproteobacteria* (*r*=0.81, *p*=0.004), *Betaproteobacteria* (*r*=0.90, *p*<0.001), *Bacteroidetes* (*r*=0.70, *p*=0.02), and *Firmicutes* (*r*=0.90, *p*<0.001). The ratios of *Alphaproteobacteria:Betaproteobacteria*, previously identified as a discriminator between Arctic and alpine cryoconite (Cook et al., 2015) were highly correlated between nanopore and pyrosequencing datasets (*r=*0.90, *p<*0.001). In summary, nanopore sequencing of 16S rRNA genes conducted according to in-field protocols correlates well with insights derived from second-generation sequencing performed in laboratory settings.

**Figure 6:**
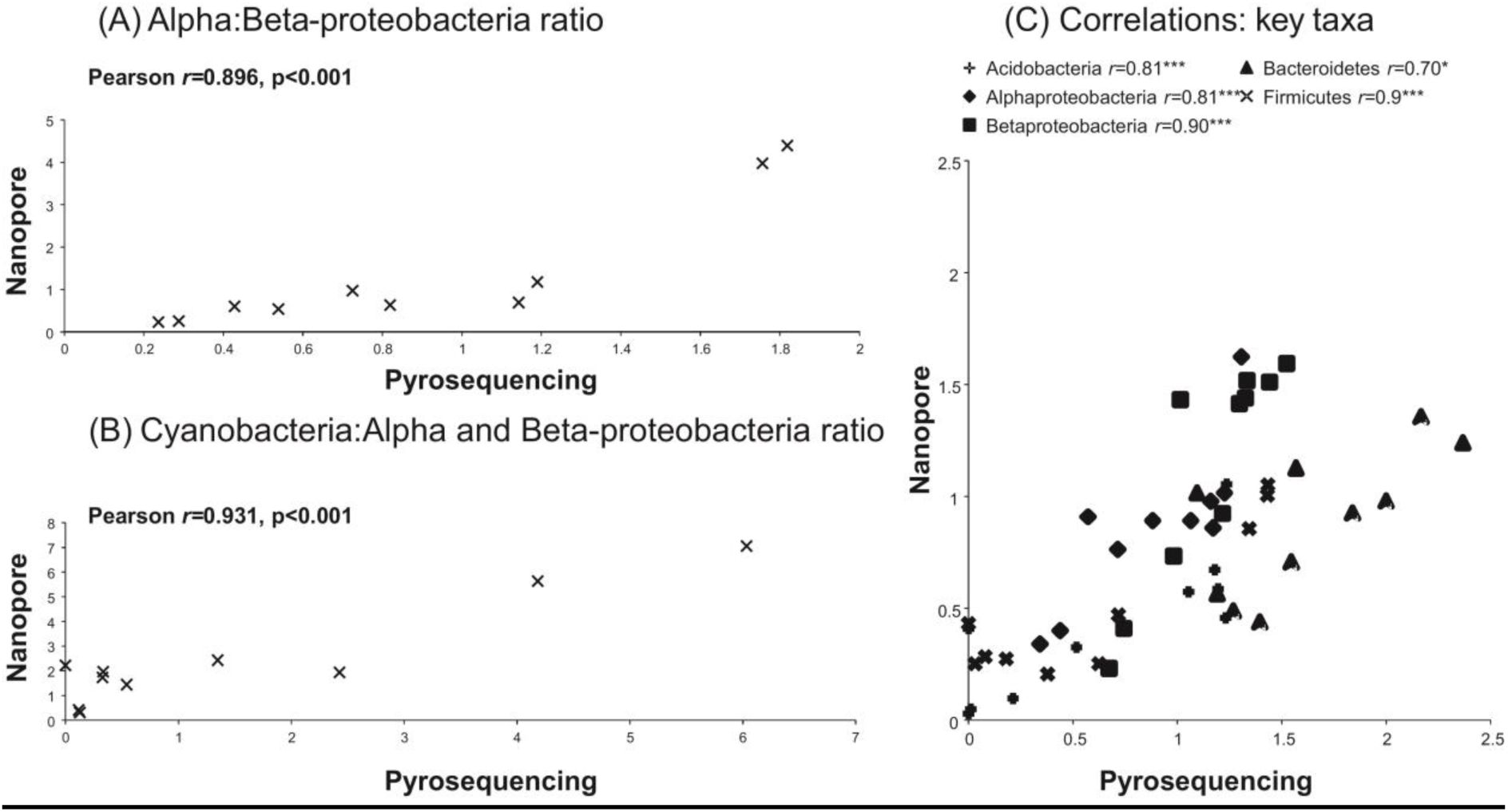
Benchmarking nanopore 16S rRNA gene amplicon sequencing of cryoconite bacterial communities by comparison with laboratory-generated data. Correlation of log relative abundances between (A) the ratio of *Alphaproteobacteria* and *Betaproteobacteria* (B) *Cyanobacteria to Alphaproteobacteria:Betaproteobacteria* and (C) key taxonomic groups revealed using nanopore and pyro-sequencing of 16S rRNA genes. Positive, significant or highly significant Pearson *r* correlations are observed for each taxon.

### Benchmarking: 16S rRNA gene nanopore sequencing of mock communities

Our in-field 16S rRNA gene analyses are limited by two caveats. Firstly, to provide rapid, in-field insights we must currently rely upon uncorrected nanopore 1D reads which are therefore subject to a relatively high error rate. In this study we anticipated the availability of full-length 16S rRNA gene reads and hence stronger phylogenetic signal relative to the random error profile of nanopore-induced error, coupled with the aggregation of reads at higher taxonomic levels would amortize the effect of diminished taxonomic resolution. Secondly, the depth of sequencing may be relatively shallow compared to laboratory-based sequencing.

Therefore, we sought to evaluate the fidelity, reproducibility and sensitivity of in-field 16S rRNA gene sequencing by using ZymoBIOMICS^™^ Microbial Community Standard mock communities where a single species, *Micrococcus luteus*, had been spiked at 16S rRNA gene concentrations in the range 0-1.2% (FIG 7). A total of 220,051 reads were de-multiplexed, with an average of 18,375 reads per sample (±1 SD=3802, range 13,165-24,858). We obtained an average of 11,728 reads per sample (±1 standard deviation: 2962) assigned to family level RDP taxonomy, of which on average 94% (±1 standard deviation: 4.8%) could be assigned to genus or *Enterobacteriaceae* level taxa.

**Figure 7:**
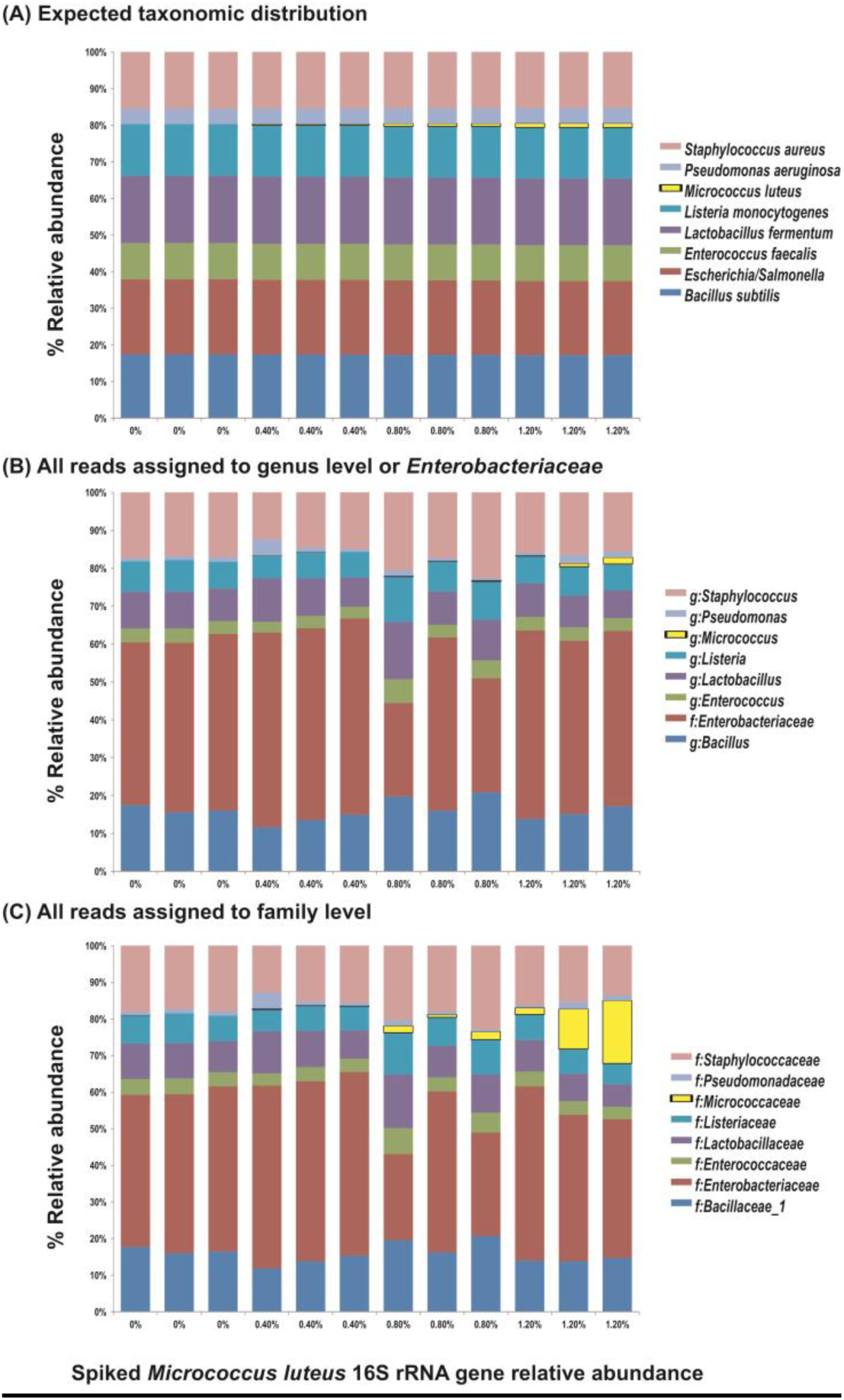
Benchmarking nanopore16S rRNA gene amplicon sequencing by sequencing mock communities using in-field protocols. Panel (A) shows the expected taxonomic distribution of The ZymoBIOMICS Microbial Community Standard for each sample, spiked in triplicate with *Micrococcus luteus* NCTC2665 genomic DNA to afford theoretical relative abundances of 16S rRNA genes in the range of 0-1.2% of the total bacterial community. Panel (B) shows shows the observed data where all reads assigned to genus level (excepting the poorly resolvable *Enterobacteriaceae* while Panel (C) shows all reads at the family level. The expected and observed *Micrococcus luteus* spike is highlighted in yellow.

The composition of the bacterial community at family or genus/*Enterobacteriaceae* level was as expected from the known composition of the mock community, and the relative abundances measured in the showed strong, significant correlations with the expected relative abundances of the ZymoBIOMICS^™^ Microbial Community Standard (family: Pearson *r=*0.74, *p=*0.035; genus/*Enterobacteriaceae*: Pearson *r=*0.73, *p=*0.038), however this was distorted by the over-representation of *Enterobacteriaceae* reads within the dataset, with very poor differentiation of *Escherichia* or *Salmonella* within the taxon. This is consistent with the weaker phylogenetic signal of 16S rRNA genes within these taxa (Fukushima et al. 2002).

To provide an insight to the sensitivity of in-field nanopore sequencing to the presence of taxa at low relative abundance, we introduced *Micrococcus luteus* NCTC 2665 genomic DNA to provide 16S rRNA gene copies to provide expected relative abundances in the range 0-1.2%. Since the ZymoBIOMICS^™^ Microbial Community Standard does not contain any *Actinobacteria*, we assumed that reads assigned to its family *Micrococcaceae* or genus *Micrococcus* corresponded to 16S rRNA genes introduced by us. In the three samples of the non-spiked baseline community, only one read was detected assigned to *Micrococcaceae*, however reads assigned to *Micrococcaceae* or *Micrococcus* were detected in two of three samples at 0.4% expected relative abundance, and then in all samples at 0.8-1.2% expected relative abundance. The relative abundance of reads assigned to *Micrococcaceae* and *Micrococcus* were positively significantly correlated with the expected relative abundance (both: Pearson *r=*0.66, *p=*0.02). While future work in developing nanopore-optimized bioinformatics pipelines for 16S rRNA gene analyses could improve reliable assignment of reads to discrete operational taxonomic units, we contend our our analyses highlight the viability of rapid, in-field sequencing of 16S rRNA genes on highly portable MinION devices for comparison of microbial communities.

### Portable microbiome sequencing: implications for microbial ecology

Within this study we describe the application of portable nanopore sequencing for the in-field characterization and comparison of microbial communities. Refinement of our approaches illustrates the potential for generating and analysing sequence data within remote locations without recourse to server-supported bioinformatics, and even in highly resource limited settings.

Advances in high throughput sequencing capacity are underpinning continual revolution, revealing revelations about the structure and potential function of microbial communities inhabiting niches throughout every conceivable habitat in the biosphere. Just over a decade ago, Curtis (2006) wrote of the urgent need for microbial ecologists to “go large” and embark upon the high throughput characterization of microbial diversity. Such data-collection initiatives represent essential pre-requisites to the development of mechanistic and predictive insights. Arguably, the vision set out by Curtis (2006) is being accomplished by initiatives such as the Earth Microbiome Project (Thompson et al., 2017), enhancing our coverage of microbial diversity across the planet. However, cataloguing microbial diversity has hitherto been contingent on laboratory-based, high-throughput sequencing platforms, replete with high levels of laboratory infrastructure. This has been at the cost of agility in characterizing and comparing microbial diversity. In-field sequencing using portable MinION sequencers and laptop-based bioinformatics approaches described herein offers the opportunity to regain agility in the characterization of Earth’s microbiomes by supporting distributed, at-source DNA sequencing. Loman and Gardy (2017) advocate the merger of human, animal and environmental genomic surveillance through scalable, portable DNA sequencing for digital epidemiology with the goal of achieving a “sequencing singularity”. Such a sequencing singularity offers benefits for navigating Earth’s microbial diversity in the broader sense also. We contend it is now time to “go small”.

“Going small” has three conspicuous advantages. Firstly, microbial processes sense and amplify the impacts of environmental change(Vincent, 2010). Understanding the genomic basis of such processes therefore represents a research priority. Parallels may be drawn with the promise of portable sequencing enabled disease diagnosis and surveillance (Gardy and Loman, 2017). For example, within the context of the global cryosphere, rapid warming is creating hitherto unseen opportunities and challenges in the study of microbial diversity. Presently, we lack DNA datasets for over 99.5%(Edwards, 2015; Edwards and Cook, 2015) of Earth’s glaciers, which represent Earth’s largest freshwater ecosystem (Edwards et al., 2014a) and yet are highly endangered by climatic warming(Hotaling et al., 2017; Milner et al., 2017). Rapid, on-site investigation of microbial interactions with climate warming has the potential to develop baseline data for fragile ecosystems such as glaciers, and to gather observational data as a prelude to hypothesis testing.

Secondly, accessing microbial diversity in remote locations, rather than merely its collection and transfer for analysis, also reduces logistical risks. This takes many forms. Sequencing on-site precludes the risk of post-collection changes in community structure, sample degradation or loss in transit *e*.*g*.(Choo et al., 2015; Hodson et al., 2017). Moreover, investigators gain the flexibility to adjust their sampling campaigns, maximising the value of field campaigns. Furthermore, retrieval of microbial diversity from inaccessible environments requires sophisticated engineering approaches to assure the integrity of recovered samples. Examples include clean subglacial lake access (Siegert et al., 2012; Christner et al., 2014). In-field sequencing offers the possibility of monitoring the recovery of high quality samples, the detection and prevention of contamination, and the optimization of sample capture events within a single field campaign.

Thirdly, in-field sequence based characterization and comparison of microbial communities offers the opportunity for distributed characterization of Earth’s microbiomes, thus expanding both our geographical and genomic coverage of microbial diveristy. Gilbert et al argue that if Darwin had the technological capacity, he would have used metagenomics in the surveys of biodiversity which underpinned the formulation of the Origin of Species(Darwin, 1859). Indeed if Darwin had been a metagenomic scientist, then *HMS Beagle* would likely have been equipped for in-field sequencing for the discovery of metagenomic diversity.

Such considerations are not whimsical. The disparity between the technological constraints on enumerating plant and animal biodiversity and microbial biodiversity underlie a schism between “animal and plant” ecology and “microbial” ecology. This is consequential in that whether microbial equivalents of long-established laws in animal and plant biogeography exist remains a contemporary research question e.g. (Carbonero et al., 2014; van der Gast, 2015). We highlight the potential for distributed discovery of Earth’s microbiomes supported by in-field DNA sequencing to underpin a new generation of scientific exploration – and explorers.

### Conclusions

Here we report the use of portable nanopore sequencing to characterize and compare microbial communities while in the field. Our approaches robustly characterize the taxonomic composition of glacial microbial communities using shotgun metagenomics, and permit their comparison by 16S rRNA gene amplicon sequencing. The experiments reported show the versatility of nanopore sequencing approaches for microbiome analyses in a range of field settings, and the coherence of data produced with established approaches for investigating microbial diversity. Continued development in the field of nanopore sequencing, for example the use of lyophilized reagents and ambient-temperature stable nanopore flow cells for sequencing with on-laptop analytic approaches raises the prospect of highly agile characterization of Earth’s microbiomes at source.

## Acknowledgements

AE is supported by a Leverhulme Research Fellowship (RF-2017-652), a Royal Geographic Society Walters Kundert Arctic Fellowship and acknowledges the support of the HEFCW Capital Research Infrastructure Fund and BBSRC funded facilities required for the analyses described herein. NERC (NE/K000942/1) supported alpine fieldwork. JMC acknowledges the Rolex Enterprise Award which supported Greenland fieldwork. ARD is supported by a Life Sciences Research Network Wales PhD scholarship while SR is supported by a Sêr Cymru Fellowship. AE, JMC and TD gratefully thank Eddie Frost and Jeffery Garriock for their support and *per*severance in the face of technical challenges on Greenland.

## Author contribution

AE: Conceived study, conducted fieldwork, sequencing, analysis, wrote manuscript. ARD: Developed DNA extraction method, conducted fieldwork. SMN, LAJM, AS and SR: bioinformatics analysis and support; BS, JMC, TD and AJH: conducted fieldwork. All authors contributed to and commented upon the manuscript.

